# A comparison of bioinformatics pipelines for compositional analysis of the human gut microbiome

**DOI:** 10.1101/2023.02.13.528280

**Authors:** Joanna Szopinska-Tokov, Mirjam Bloemendaal, Jos Boekhorst, Gerben DA Hermes, Thomas HA Ederveen, Priscilla Vlaming, Jan K Buitelaar, Barbara Franke, Alejandro Arias-Vasquez

## Abstract

Investigating the impact of gut microbiome on human health is a rapidly growing area of research. A significant limiting factor in the progress in this field is the lack of consistency between study results, which hampers the correct biological interpretation of findings. One of the reasons is variation of the applied bioinformatics analysis pipelines. This study aimed to compare five frequently used bioinformatics pipelines (NG-Tax 1.0, NG-Tax 2.0, QIIME, QIIME2 and mothur) for the analysis of 16S rRNA marker gene sequencing data and determine whether and how the analytical methods affect the downstream statistical analysis results. Based on publicly available case-control analysis of ADHD and two mock communities, we show that the choice of bioinformatic pipeline does not only impact the analysis of 16S rRNA gene sequencing data but consequently also the downstream association results. The differences were observed in obtained number of ASVs/OTUs (range: 1,958 - 20,140), number of unclassified ASVs/OTUs (range: 210 - 8,092) or number of genera (range: 176 - 343). Also, the case versus control comparison resulted in different results across the pipelines. Based on our results we recommend: i) QIIME1 and mothur when interested in rare and/or low-abundant taxa, ii) NG-Tax1 or NG-Tax2 when favouring stringent artefact filtering, iii) QIIME2 for a balance between two abovementioned points, and iv) to use at least two pipelines to assess robustness of the results. This work illustrates the strengths and limitations of frequently used microbial bioinformatics pipelines in the context of biological conclusions of case-control comparisons. With this, we hope to contribute to “best practice” approaches for microbiome analysis, promoting methodological consistency and replication of microbial findings.

**Author Summary:** Studies increasingly demonstrate the relevance of gut microbiota in understanding human health and disease. However, the lack of consistency between study results is a significant limiting factor of progress in this field. The reasons for this include variation in study design, sample size, bacterial DNA extraction and sequencing method, bioinformatics analysis pipeline and statistical analysis methodology. This paper focuses on the variation generated by bioinformatics pipelines. A choice of a bioinformatic pipeline can influence the assessment of microbial diversity. However, it is unclear to what extent and how the results and conclusion of a case-control study can be influenced. Therefore, we compared the results of a case-control study across different pipelines (applying default settings) while using the same dataset. Our results indicate a lack of consistency across the pipelines. We show that the choice of bioinformatic pipeline not only affects the analysis results of 16S rRNA gene sequencing data from the gut microbiome, but also the associated conclusions for the case-control study. This means different conclusions would be drawn from the same data analysed with different bioinformatic pipeline.

## 1. Introduction

Investigations of the role of the human gut microbiota have attracted much attention in the last 15 years [1]. Specifically, results of studies of the 16S rRNA marker gene (16S) have been crucial in understanding the role the gut microbiota play in multiple common diseases, such as irritable bowel syndrome [2], autism [3], depression [4] or attention deficit hyperactivity disorder (ADHD) [5]. Although a few papers suggested best practice for microbiome analysis [6, 7], still there is a broad choice in microbiome methods. This affects the consistency across the studies. So far, 16S rRNA gene sequencing is one of the most commonly used method to study bacterial phylogeny and genus/species classification [8]. 16S rRNA gene sequencing is used as a tool to identify multiple bacterial taxa and assist with differentiating between closely related bacteria.

The classification of microbial taxonomy using the 16S rRNA gene is influenced by several factors, ranging from study design, sample size, the choice of variable region of 16S rRNA gene to sequence [9], collection and storage procedure, wet lab approaches, such as DNA extraction [10], sequencing technique and bioinformatic pipeline settings, such as frequency filters, and the taxonomic classification database [11]. Bioinformatics pipelines differ in approaches, such as quality control and filtering of the sequenced data (i.e., chimera detection, filtering sequences, denoising), Operational Taxonomic Units (OTUs) clustering algorithms or Amplicon Sequence Variant (ASV), and statistical analysis (when a statistical analysis step is included in the pipeline). All these choices may result in differences in the (observed) distribution of taxonomic groups, significantly affecting the putative relationships between the gut microbiota and disease outcomes. This limits the precision of biological and statistical conclusions, resulting in a lack of consistency between studies [5, 8, 9, 12].

In this paper, we focused on comparing bioinformatics pipelines, as their contribution to biological conclusions of microbiome studies is not sufficiently quantified. So far, studies investigating differences between bioinformatics pipelines have focused on general characteristics of the OTUs/ASVs/reads, such as richness, diversity and microbial compositional profiles, rather than on the biological conclusions that could be drawn from analyzing these characteristics [6, 13, 14]. Recently, Ducarmon et al. (2020) showed that the NG-Tax 1.0 [15] and QIIME2 [16] bioinformatics pipelines performed equally well in terms of microbial diversity and compositional profiles for 24 samples across eight types of biological specimens from human niches [13]. Poncheewin et al. (2020) compared NG-Tax 2.0 with QIIME2 using 14 mock community samples [17]. Precision of NG-Tax 2.0 (0.95) was significantly higher compared to QIIME2 (0.58). Prodan et al. (2020) used a large dataset of 2,170 samples and one mock community of 16S rRNA data to compare QIIME-uclust [18], mothur [19], USEARCH-UPARSE [20], DADA2 [21], QIIME2-Deblur [16, 22] and USEARCH-UNOISE3 [23] pipelines, and concluded that *“DADA2 is the best choice for studies requiring the highest possible biological resolution (e.g. studies focused on differentiating closely related strains)”* [6]. López-García et al. (2018) showed that when the SILVA reference database was used in combination with QIIME [24] or mothur [19] pipelines, richness and composition of 18 samples were highly similar [14]. However, beta-diversity clustered by pipelines, which they attributed to differences in less abundant bacteria. While this was not tested by López-García et al., this description hints at the possibility of different biological conclusions depending on a choice of pipeline. Only one study, Allali et al. (2017), investigated whether the same biological conclusion was reached when using different pipelines based on 14 chicken cecum 16S rRNA samples across three different treatment groups. They tested different settings of QIIME1, UPARSE and DADA2 and concluded that, despite differences in diversity and abundance, they could discriminate samples by treatment, leading to similar biological conclusions [25]. This conclusion was limited to beta-diversity (global microbiome community differences), not including a comparison of individual genera. As they reported differences in relative abundances of specific genera between pipelines, their data suggests that different pipelines could result in different lists of genera being significantly associated with a treatment.

While the existing comparisons have been essential for the field, they fall short in contributing highly-needed conclusions on how the choice of bioinformatic pipeline affects downstream statistical comparisons of microbial composition of groups (for example, humans with and without a disease). Such comparisons are also vital for the growth and stability of the field [12]. Moreover, frequently used pipelines, NG-Tax1, NG-Tax2, QIIME1, QIIME2 and mothur, have not yet been compared using the same dataset. Based on these gaps and limitations in the state of the art of the field, we aimed to determine the differences in taxonomic output between these five pipelines and how such differences affect downstream statistical analyses and interpretation of the observed results. We used the V4 16S rRNA gene sequencing data of a human case-control study of attention-deficit/hyperactivity disorder (ADHD) as well as two mock communities. We would like to highlight that our aim is not to draw biological conclusions from these analyses (for this we refer to [26]), but rather highlight differences brought in by the choice of bioinformatic pipeline.

## 2. Materials and Methods

### 2.1. Dataset

The material and methods and the results sections are divided into two parts: (i) results based on clinical samples (NeuroIMAGE dataset [26]) and (ii) results based on mock communities (MC), which allow us to better interpret the results based on the clinical samples.

#### 2.1.1. NeuroIMAGE dataset

We used the clinical and microbial information from a group of samples belonging to a case-control sample (case, n=42; control, n=50) reported in the NeuroIMAGE study [26]. For an exhaustive description of the sample, inclusion criteria, ADHD analysis methods, diagnostic procedures, and study design used in this study, see Szopinska-Tokov et al., 2021 [26], of which this study is an extension.

#### 2.1.2. Mock communities

In addition to the case-control dataset, we analyzed eight samples based on two defined Mock Communities (MCs; MC3, n=4; MC4, n=4), of which one (MC4) included taxa with very low abundances (0.1%, 0.01% and 0.001%). Both MCs included the same 36 genera, but in different distributions. The laboratory processing and evaluation of the observed MC composition was done exactly the same as for the clinical samples [26]. The laboratory processing and evaluation of the expected microbial communities’ composition was carried out as described previously [15]. In short, the bacteria were grown as pure cultures and their DNA was then mixed in specific amounts for each community (the process was carried at the Laboratory of Microbiology, Wageningen University, The Netherlands). The bacterial composition of the MCs was determined with HiSeq2000, and for each bacterium used in the MCs, the full length 16S gene was sequenced with Sanger sequencing to confirm their identity.

### 2.2. Bioinformatics pipelines and their evaluation

We investigated five different pipelines: both versions of the NG-Tax pipeline (NG-Tax v.1.0 [15] and v.2.0 [17], here named NG-Tax1 and NG-Tax2), adapted QIIME (v.1.8.0; here called QIIME1) [18], QIIME2-DADA2 (v.2019.7.0; here called QIIME2) [16], and mothur (v.1.43.0) [19]. NG-Tax1, NG-Tax2 and QIIME2 are ASV-based methods, whereas QIIME1 and mothur are OTU-based methods.

The bioinformatic pipeline evaluation involved two steps: (i) bioinformatical processing and (ii) statistical testing, involving data analysis and quantification (Figure 1).

**Figure 1.**
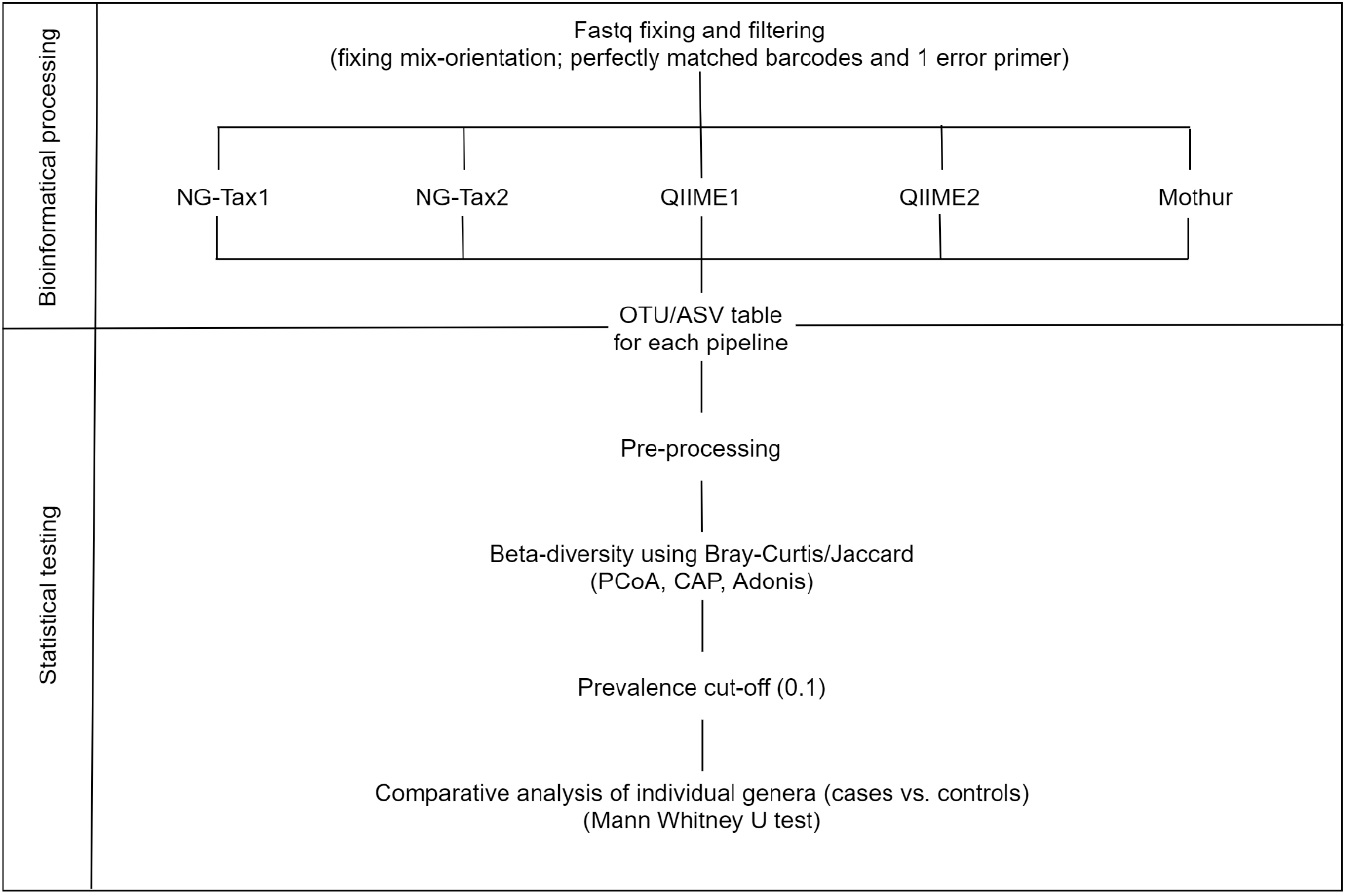
Overview of the bioinformatical and statistical steps used in this study. Top panel: Raw sequencing data (paired-end fastq file) was pre-processed; Reads were put in the same orientation. Subsequently, read pairs with perfectly matching (forward and reverse) barcodes and a maximum of one nucleotide mismatch for each (forward and reverse) primer were included in further steps. This was used as input for all pipelines (see Methods section). This resulted in the OTU/ASV tables (one for each pipeline) which were then subjected to pre-processing. Bottom panel: all statistical tests were carried out separately for each pipeline, except for beta-diversity where OTU/ASV tables were merged to directly compare the taxonomy tables between the pipelines. Prior to comparative analysis the prevalence cut-off was applied (for more details see Discussion section). For details for each step please see the main text.

#### 2.2.1. Bioinformatical processing

Before applying the pipelines, we applied an in-house script to make sure that the input was the same for all the pipelines. First, we had to deal with the mixed orientation of the sequences. This means that forward and reverse files contained both forward and reverse sequences. NG-Tax 1 and NG-Tax 2 deal with this as a part of the default settings, but this is not so straightforward for other pipelines. Second, not every pipeline can deal or deals in the same way with dual barcodes. Third, different primer settings are applied by each pipeline. In order to eliminate pipeline bias related to primer and barcode mismatch, we applied the same settings for all the pipelines. The output of the in-house script resulted in fixed orientation of the sequences having perfectly matching forward and reverse barcodes with only one nucleotide mismatch allowed for each (forward and reverse) primer. This was used as an input for all the pipelines. Furthermore, we used the default setting of the pipelines, except for taxonomic database where we used SILVA (v.132) database for all pipelines, changing the default option for NG-Tax1 and QIIME1. We used the Galaxy platform to run NG-Tax1 and NG-Tax2 (http://wurssb.gitlab.io/ngtax/galaxy.html). QIIME1 was run according to the in-house (NIZO, Ede, The Netherlands) protocol as described previously [10]. For QIIME2, we followed the “Moving Pictures” tutorial (https://docs.qiime2.org/2019.4/tutorials/moving-pictures/), and for mothur the “MiSeq SOP” (https://mothur.org/wiki/miseq_sop/).

#### 2.2.2. Statistical testing

##### 2.2.2.1. NeuroIMAGE dataset

###### 2.2.2.1.1. Pre-processing

Taxonomical names were formatted across the pipelines, e.g., D_0_Bacteria was changed into Bacteria in order to align the format of taxonomic names across the pipelines. The original sample contained a subthreshold-ADHD group [26], which was removed in the current analysis. Furthermore, we determined a threshold of total read counts based on rarefaction plots (data not shown), in order to exclude samples with small number of total reads while keeping the maximum number of samples (as explained in the ‘Moving pictures’ QIIME2 tutorial [28]). Thus, samples below 1000 total reads were not included in further analysis; this resulted in removal of two samples across all pipelines, which had on average 11 (range: 4-21) and 255 (range: 150-341) total read counts across the pipelines. The final dataset included 40 cases and 50 controls.

###### 2.2.2.1.2. OTU/ASV/reads table characteristics

As a first part of the analysis, we compared the results of the pipelines in terms of characteristics and distribution of reads, OTUs/ASVs, singletons (a single sequence), unclassified reads, and taxa. The analyses were focused on the genus level, since this is the level at which most (clinical) studies focus to identify an association with a disease/disorder status. This is due to the fact that analysis based on 16S rRNA gene hypervariable region(s) limits the taxonomic resolution to family- or genus-level [29].

We visualized overlapping genera between the pipelines using a Venn Diagram. In order to see how the percentage of overlapping genera changed based on different filtering thresholds, we compared the gut microbiome composition of: A) all the genera, B) genera after applying a 10% prevalence cut-off, C) genera with relative abundance >0.1%, and D) genera with relative abundance <0.1%.

###### 2.2.2.1.3. Beta-diversity

While beta diversity analysis is typically performed at the level of OTU/ASV, we did it at the genus level in order to be able to compare the microbial composition (relative abundance; Bray-Curtis dissimilarity metric) and structure (presence/absence; Jaccard similarity index) [30] across different bioinformatics pipelines. The statistical significance of this comparison was determined using Permutational Multivariate Analysis of Variance (PERMANOVA) using the R package ‘adonis’ for all pipelines; as a post hoc analysis, we performed pairwise analysis between all pipelines [31]. The results were visualized by unconstrained (Principal Coordinate Analysis, PCoA) and constrained (Canonical Analysis of Principal coordinates, CAP) ordination methods [32] by applying following formula: ordinate(ps.merged.rel, “CAP”, “bray”, ^~^ Pipeline). Additionally, we computed Tukey Honest Significant Differences (TukeyHSD; calculated based on betadisper using the R package ‘vegan’ [31, 33, 34]) to expand the PCoA analysis and to investigate intra-sample variation in a pairwise comparison manner.

###### 2.2.2.1.4. Comparative analysis at the genus level

In order to obtain a more detailed overview of microbiome composition differences, we compared the pipelines (i) in terms of the relative abundance of the ten most abundant genera (in order to maximize our ability to find differences between the groups) and (ii) between cases and controls. At this stage, we filtered out unclassified genera and applied a prevalence cut-off of 10% (at the genus level), meaning that only genera present in >10% of the total number of samples were kept, in order to keep the most informative data for the downstream statistical analysis [26]. Next, given the zero-inflated nature of the data, a non-parametric (rank-based) test (Mann-Whitney U) was applied to evaluate significant differences in relative abundances of bacterial genera between cases and controls. As we aimed to evaluate the effects of the different pipelines rather than scale and significance of the differences between them, this method seemed appropriate (see [12] for an extensive comparison of abundance testing methods).

In analysing the consistency pattern of the case-control association results across pipelines, we assigned a bioinformatics pipelines P-value Consistency Score (PCS, ranging from zero to five) to score the number of pipelines showing statistically significant differences between groups per each genus (P<0.05 unadjusted). A PCS=5 meant that all pipelines found significant differences (P<0.05 unadjusted) between cases and controls for a particular taxonomic group. Additionally, we calculated a genus relative abundances case/control ratio (called Fold-Change, FC) and compared it (as an effect measure) between the pipelines. The FC was calculated by using the foldchange() function from the “gtools” package (v.3.8.1) [35]. FC was computed as follows: case/control if case>control, and as - control/case otherwise. Furthermore, we tested the correlation between the PCS and the average relative abundance (RA; per genus for all the pipelines) and average percentage of zeros of each genus based on all pipelines.

All analyses were performed in RStudio (v.1.2.5033; R v.3.6.3) [36] using “phyloseq” (v.1.28.0) [37], “microbiome” (v.1.6.0) [38], and “vegan” R packages [34], visualized by using “ggplot2” [39] (v.3.3.0), “VennDiagram” [40] (v.1.6.20), “ggpubr” [41] (v.0.2.4), and “heatmaply” [42] (v.1.1.0) R packages; statistical analyses where performed by using the “stats” R package (v.3.6.3) [39].

##### 2.2.2.2. Mock communities

The main focus of the MC analysis was to compare observed to expected MC composition in order to further evaluate the reliability and comparability of the pipelines. First, we compared the number of genera observed to the expected MC composition. Second, beta-diversity was analysed as described above. Third, we calculated Spearman’s rho statistic via “stats” R package (v.3.6.3) [39] to (i) compare the observed to the expected MC composition (relative abundance), and to (ii) compare the pipelines against each other. In this way, we could identify the strength of correlation between the pipelines, and identify strength of correlation between the pipelines and the expected MC composition. The results were visualized by a heatmap using the “heatmaply” (v.1.1.0) R package [42] to identify any inconsistencies across the pipelines.

## 3. Results

### 3.1. NeuroIMAGE dataset

#### 3.1.1. OTU/ASV/reads table characteristics

Table 1 shows the characteristics and distribution of OTUs/ASVs/reads per bioinformatic pipeline for the complete study (N=90). We observed a high degree of variation across the pipelines for all the variables. The total number of reads varied across the pipelines with QIIME1 showing the highest and QIIME2 the lowest number of reads (percentage difference = 38.2%). Moreover, QIIME1 and mothur showed the highest number of OTUs/ASVs, NG-Tax1 and NG-Tax2 showed the lowest (relative difference ranging from 77.9% to 164.6%). Mothur showed the highest number of singletons (69.2% of the total OTUs), but these only accounted for 0.67% of the total reads; these singletons did not influence significantly the total relative abundance (when singletons were removed, the relative abundance of other taxa was not influenced, data not shown). Furthermore, mothur and QIIME1 detected the biggest percentage of unclassified OTUs/ASVs (46.1% and 40.2%, respectively, at the genus level), QIIME2 the lowest (4.7%).

**Table 1.** Summary of OTU/ASV characteristics between bioinformatics pipelines.

|  | NG-Tax1 | NG-Tax2 | QIIME1 | QIIME2 | mothur |
| --- | --- | --- | --- | --- | --- |
| Total number of final reads | 1,414,916 | 1,357,891 | 1,692,581 | 1,149,886 | 1,390,041 |
| Median of final reads per sample (IQR) | 14,619<br>(7,648-20,997) | 13,925<br>(7,411-19,998) | 17,315<br>(8,783-25,819) | 11,519<br>(5,385-17,515) | 14,200<br>(7,173-20,742) |
| Total number of identified OTUs/ASVs | 1,958 | 1,958 | 20,140 | 4,458 | 13,392 |
| Number of singletons (% of total number of OTUs/ASVs) | 0 | 0 | 1,291 (6.41) | 3 (0.07) | 9,269 (69.21) |
| Number of singletons (% of total number of OTUs/ASVs) before pre-processing step | 0 | 0 | 0 | 7 (0.14) | 10,206 (69.50) |
| Number of unclassified reads at the genus level (% of total reads) | 202,165 (14.3) | 193,698 (14.3) | 427,601 (25.3) | 23,091 (2.0) | 23,404 (1.7) |
| Number of unclassified OTUs/ASVs at the genus level (% of total number of OTUs/ASVs) | 321 (16.4) | 321 (16.4) | 8,092 (40.2) | 210 (4.7) | 6,170 (46.1) |
| Number of genera | 177 | 176 | 312 | 254 | 343 |
| Number of genera remaining after using a prevalence cut-off of 10% (% of total genera) | 74 (41.8) | 74 (42.1) | 145 (46.5) | 115 (45.3) | 142 (41.4) |
| Number of genera below 0.1% relative abundance (% of total genera) | 115 (65) | 115 (65.3) | 243 (77.8) | 186 (73.2) | 275 (80.2) |
| Number of phyla | 10 | 10 | 13 | 14 | 15 |

IQR = interquartile range
Important to mention, the number of singletons for QIIME1 was the effect of pre-processing (removal of the subthreshold group and samples having > 1000 reads). As a default setting, all the pipelines, except QIIME2 and mothur, remove singletons (see Number of singletons (% of total number of OTUs/ASVs) before pre-processing step).

Of the genera detected by NG-Tax1, NG-Tax2, QIIME1, QIIME2 and mothur, only 40% overlapped between all pipelines (Figure 2A). After applying the 10% prevalence cut-off to preserve the most informative data for the downstream statistical analysis, 41.4% to 46.5% of the genera remained (Table 1). The prevalence cut-off did not improve the percentage of overlapping genera (Figure 2B), indicating that more prevalent genera are not necessarily shared across the results from the different pipelines. The relative abundance threshold did improve the percentage of overlapping genera; genera above 0.1% were more commonly shared across pipelines (70%) than genera below 0.1% (20%) (Figure 2C,D).

**Figure 2.**
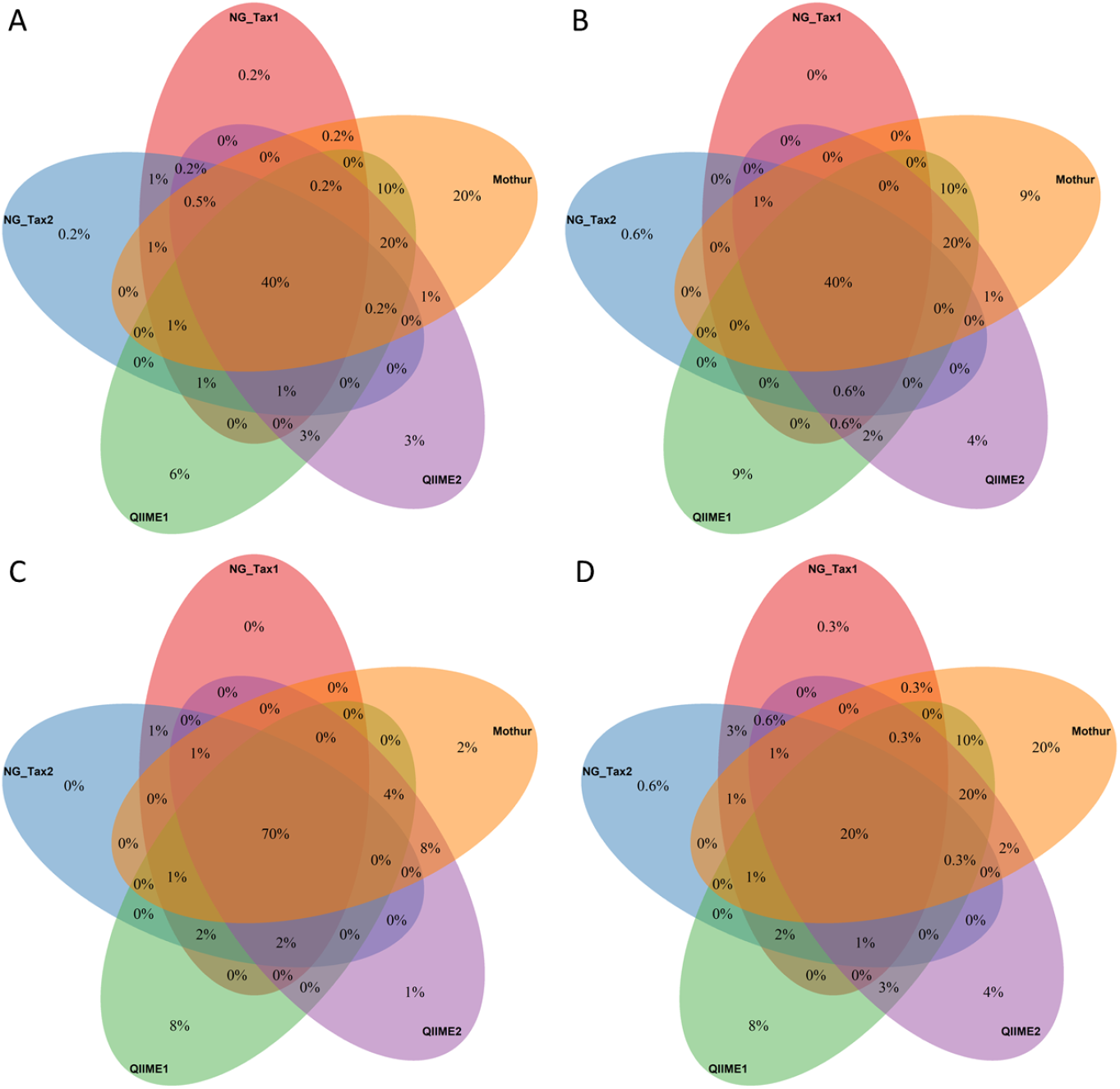
Venn diagram showing overlap between genera produced by five different bioinformatics pipelines. A) represents the overlap of genera based on raw data (based on 413 genera across pipelines), B) represents the overlap of genera after a 10% prevalence cut-off across samples (based on 171 genera across pipelines), C) overlap of genera with relative abundance >0.1% (N=80, genera across pipelines), and D) overlap of genera with relative abundance <0.1% (N=357 genera across pipelines).

#### 3.1.2. Beta-diversity

Unconstrained PCoA plots based on the Bray-Curtis measure revealed that samples clustered based on the sample ID rather than the bioinformatics pipelines (Figure 3A). However, the constrained ordination method, CAP analysis, indicated relevant differences between the pipelines in terms of microbial composition (Bray-Curtis index) at the genus level (Figure 3B). The CAP analyses captured the variation in community structure in the first two components (CAP 1 and CAP 2) accounting for 11.1% of the total variance (Figure 3B). The same results were observed in terms of microbiome structure using Jaccard’s similarity index (Figure S1). PERMANOVA analysis supported the results by revealing that microbial composition (Bray-Curtis: R^2^=13.9%, p<0.001) and structure (Jaccard: R^2^=9.5%, p<0.001) differed significantly between the pipelines and, as expected, more variability was explained by the same sample ID (Bray-Curtis: R^2^=89.5%; p<0.001 and Jaccard: R^2^=82.8%; p<0.001). Additionally, we performed a pairwise comparison of group means dispersions (TukeyHSD). The analysis confirmed that the intra-sample variation is quite similar across the pipelines, except for QIIME1 (Figure 3C).

**Figure 3.**
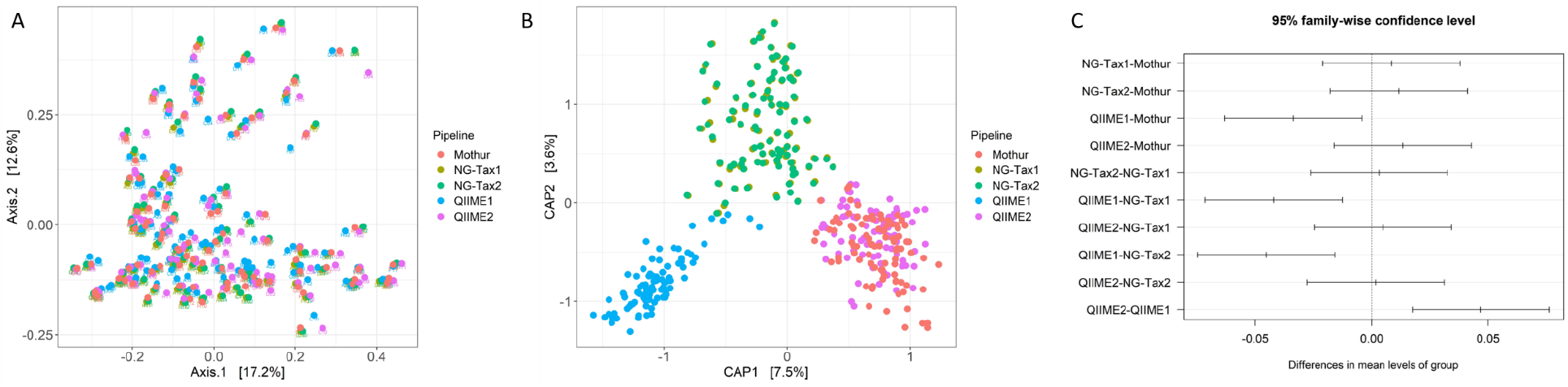
Results for the Bray-Curtis dissimilarity metric. A) Principal Coordinates Analysis (PCoA) plots with the percentage explained variance by the principal coordinates. B) Canonical Analysis of Principal coordinates (CAP) ordination plot of structure in microbial communities associated with bioinformatics pipelines. C) TukeyHSD, a pairwise comparison of group mean dispersions revealed that the intra-sample variation was quite similar across pipelines, with QIIME1 forming the exception.

The CAP analysis also showed that NG-Tax1 and NG-Tax2 clustered together, and QIIME2 clustered with mothur (Figure 3C,D). We investigated these results in more detail, by running PERMANOVA again, this time only with NG-Tax1 and NG-Tax2 or with QIIME2 and mothur, to investigate how statistically different these clusters were. The results indicated statistically significant differences between the pipelines, however, with very small percentages of explained variation (NG-Tax1/NG-tax2 R^2^=0.016%, p<0.001; QIIME2/mothur R^2^=0.9%, p<0.001; the results of pairwise PERMANOVA analyses for other combinations can be found in Supplementary Table S1).

#### 3.1.3. Comparative analysis of individual genera

We also compared the distribution of the ten most abundant genera found by each pipeline (Figure 4). These genera were not identical across the pipelines: across the five pipelines, 16 unique genera were observed. The RA values for all of the 16 unique genera were statistically significantly different between pipelines (Friedman test, Bonferroni-adjusted p-values <0.001). The descriptive statistics of this data can be found in Supplementary Table S2.

**Figure4.**
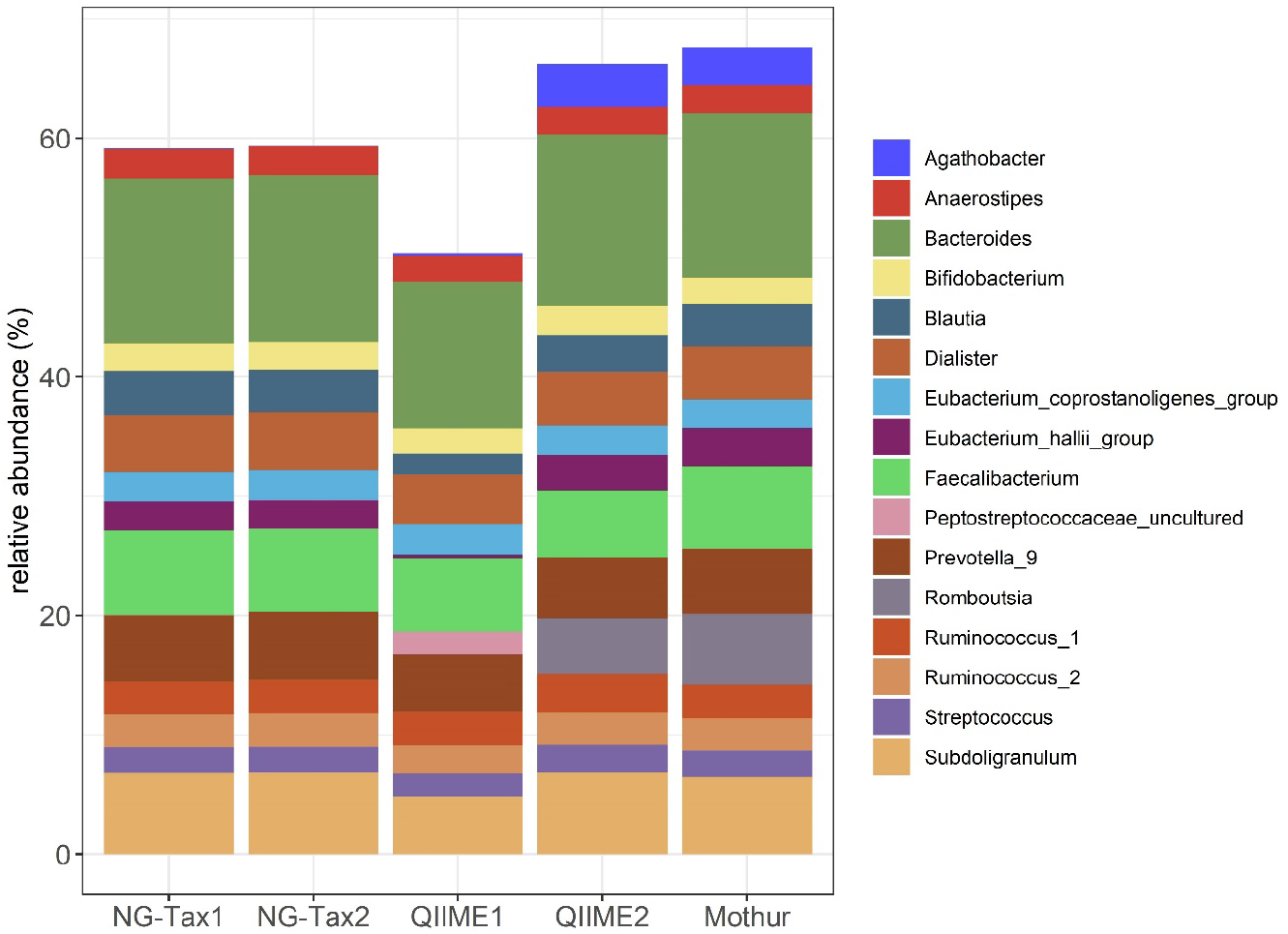
Bacterial genera profile. Top 10 most abundant bacterial genera per pipeline resulted in a total of 16 unique genera. We excluded unclassified genera, since they represent a group of genera rather a single genus.

#### 3.1.4. Taxonomic differences between cases and controls across pipelines

We carried out univariate testing of the relative abundance of individual genera between ADHD cases (N=40) and controls (N=50) in order to investigate if the downstream statistical conclusions were consistent across the pipelines. In total, 10 genera showed nominally significant differences (p< 0.05) between cases and controls in at least one pipeline (Table 2), but these differences were not consistent across all pipelines. Based on the P-value consistency score (PCS), only one of the 10 genera showed total agreement in terms of PCS (PCS=5), none showed high agreement (PCS=4), three genera showed moderate agreement (PCS=3), and two genera showed partial agreement (PCS=2). The rest of the genera (N=4) showed no agreement (PCS=1) (Table 2). The descriptive statistics of the 10 genera can be found in the Supplementary Table S3.

**Table 2.** The table represent a fold change (case/control ratio), p-value consistency score (PCS), and percentage of zeros for genera which were nominally significant (p<0.05) different between cases and controls by at least one pipeline. Values highlighted in red indicate nominal significance (p<0.05). A negative value indicates that the cases’ mean is lower than the controls’ mean.

| Genera | NG-Tax1 | NG-Tax2 | QIIME1 | QIIME2 | mothur | PCS | % of zeros |
| --- | --- | --- | --- | --- | --- | --- | --- |
|  | Fold Change |  |  |  |  |  |  |
| Coprococcus_2 | 1.09 | 1.12 | -1.24 | -1.06 | 1.05 | 5 | 40 |
| Prevotella_9 | -1.83 | -1.81 | -2.02 | -1.76 | -1.87 | 3 | 35 |
| Ruminococcus_1 | -1.51 | -1.50 | -1.49 | -1.49 | -1.55 | 3 | 8 |
| Eubacterium_eligens_group | -1.61 | -1.48 | -1.58 | -1.92 | -1.65 | 3 | 62 |
| Tyzzarella_3 | 1.02 | -1.02 | NA | 1.88 | 1.77 | 2 | 74 |
| Howardella | NA | NA | 4.45 | 4.88 | NA | 2 | 82 |
| Eubacterium_ventriosum_group | -2.32 | -2.34 | -1.93 | -2.31 | -2.02 | 1 | 17 |
| Fusicatenibacter | -1.66 | -1.69 | 1.10 | 1.24 | 1.03 | 1 | 47 |
| Clostridiales_vadinBB60_group_uncultured_bacterium | NA | NA | 1.19 | 2.97 | NA | 1 | 74 |
| Lachnospiraceae_UCG_004 | NA | NA | -1.56 | NA | -1.13 | 1 | 49 |

In order to determine the effect of the differences in genus abundance on the case-control comparison between the pipelines, we compared Fold Change (FC) based on genera relative abundance (Table 2). Three observations stand out. First, the FC differs between the pipelines. For example, for *Clostridiales_vadinBB60_group_uncultured_bacterium*, QIIME1 resulted in a case/control ratio of 1.19, whereas QIIME2 resulted in a ratio of 2.97. Second, for both versions of QIIME, the FC of *Coprococcus_2* was in the opposite direction compared to the other three pipelines. Third, in some cases (e.g., *Prevotella_9*, *Ruminococcus_1*), the FC was almost the same between the pipelines, but still only one pipeline indicated nominal significance.

In general, some genera were missing in some pipelines, and there were differences in effect size or even in direction between pipelines for genera that were nominally significant different between cases and controls. The non-parametric rank test indicated that genera present in all pipelines (N=6) differed statistically in their relative abundance among the pipelines (Friedman test, Bonferroni-adjusted p-values <0.002, Supplementary Table S3).

Testing the correlation between PCS and two measures of frequency, relative abundance and the percentage of zeros, we found the correlation coefficient between PCS and relative abundance to be r_PCS-RA_=0.58 and the one between PCS and percentage of zeros to be r_PCS-%0_=−0.24 (Figure S2A,B). Both correlations were non-significant (p>0.05), however, suggesting that the consistency across the pipelines was independent of bacterial relative abundance and the observed percentage of zeros. The lack of significance should be treated with caution, as it could be a result of the low number of features included in the analysis (n=10 genera).

### 3.2. Mock communities

#### 3.2.1. Genus richness

Mothur identified the highest and NG-Tax1 and NG-Tax2 the lowest number of genera in both MCs. NG-Tax1 (N_MC3_=31, N_MC4_=25), NG-Tax2 (N_MC3_=31, N_MC4_=25) and QIIME2 (N_MC3_=39, N_MC4_=36) approached the expected genus richness (N_MC3_=36, N_MC4_=36) closer than QIIME1 (N_MC3_=64, N_MC4_=67) and mothur (N_MC3_=84, N_MC4_=101) (Table S4).

#### 3.2.2. Beta-diversity

We also compared the observed and expected beta-diversity (at genus level) in the MCs. PCoA plots based on Bray-Curtis and Jaccard measures revealed that samples clustered based on the pipelines (Figure 5). 90% (for MC3) and 98% (for MC4) of total microbial composition variance (Bray-Curtis, p_MC3_<0.001 and p_MC4_<0.001) and 87% (in case of MC3) and 97% (in case of MC4) of total microbial structure variance was explained by pipelines (Jaccard, p_MC3_<0.001 and p_MC4_<0.001).

**Figure 5.**
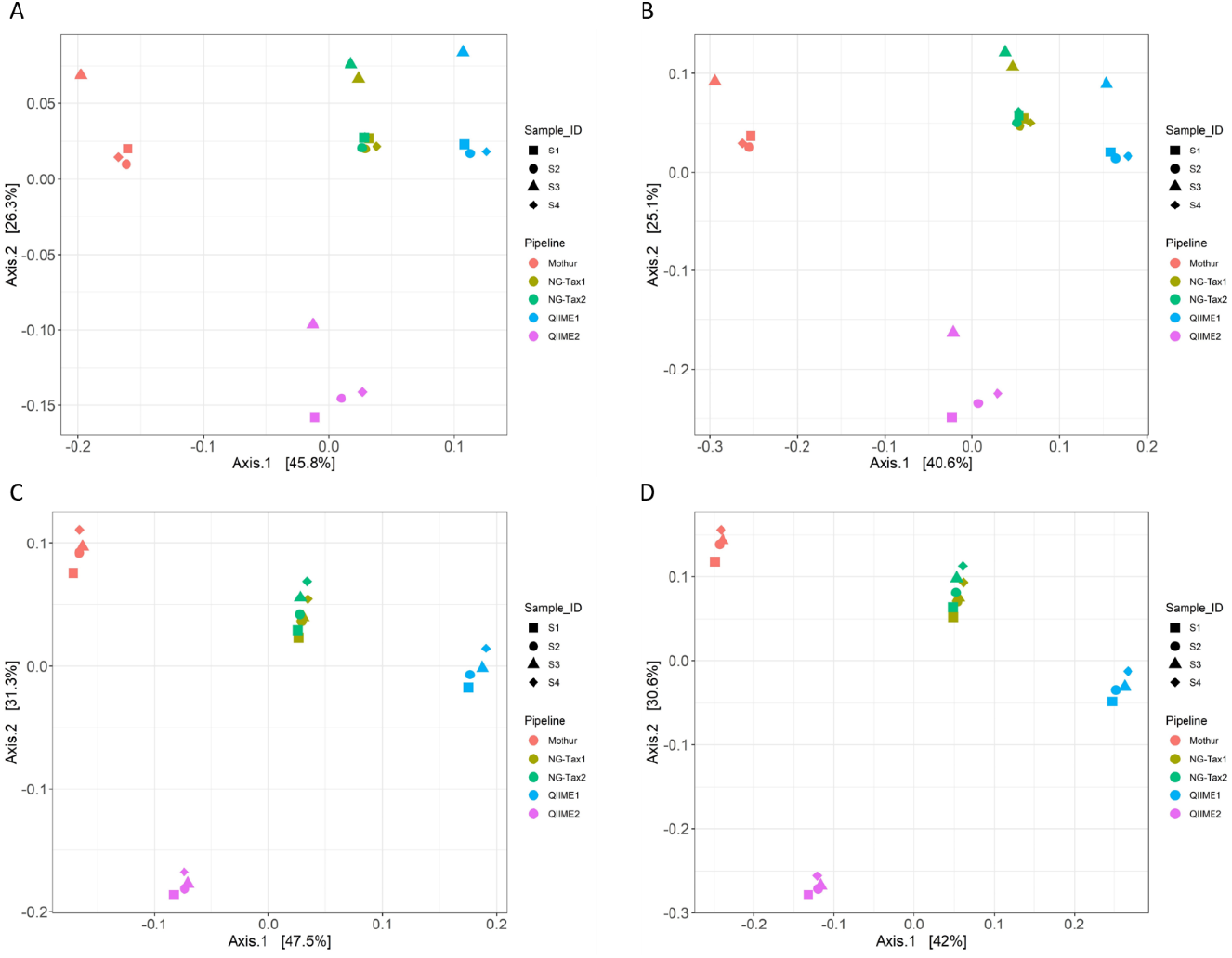
PCoA of MC composition was affected by the choice of bioinformatics pipelines. Results of the Bray-Curtis dissimilarity metric and Jaccard similarity index based on MC3 are shown in panel A and B, respectively, and based on MC4 are shown in C and D, respectively. S1 = Sample 1.

#### 3.2.3. Correlation analysis

The correlation of observed and expected MC relative abundance (based on N=36 genera) showed that

QIIME2 had the highest correlation coefficient (r_MC3_=0.70, r_MC4_=0.76), followed by mothur (r_MC3_=0.67, r_MC4_=0.65), QIIME1 (r_MC3_=0.61, r_MC4_=0.64), NG-Tax1 (r_MC3_=0.56, R_MC4_=0.61) and NG-Tax2 (r_MC3_=0.56, r_MC4_=0.61) (Figure S3 A,B).

#### 3.2.4. Comparative analysis of individual genera

Comparison of individual genera showed inconsistencies across pipelines for both MCs (Figure S4, S5). For example, NG-Tax1 and NG-Tax2 did not detect *Enterobacter* and *Dorea*, while QIIME2 did not detect *Serratia*, mothur did not detect *Klebsiella*, while QIIME1 did not detect *Anaerostipes* from either MCs. All pipelines failed to classify *Salmonella.* Some pipelines under/overrepresented certain genera; for example, QIIME1 overrepresented *Enterobacter* and *Pseudomonas;* NG-Tax1 and NG-Tax2 overrepresented *Klebsiella*. As expected, NG-Tax1 and NG-Tax2 did not detect genera below 0.1% abundance included in MC4 (due to the abundance cut-off setting) (Figure S5), whereases QIIME2 did not detect genera below 0.01%.

## 4. Discussion

### Summary

In this study, we compared five frequently used bioinformatics pipelines for the processing of 16S rRNA gene amplicon sequencing data, NG-Tax1, NG-Tax2, QIIME1, QIIME2 and mothur, to determine whether and in which way the analytical methods of each of these pipelines affect the downstream statistical analysis results. For this purpose, we used a clinical (case-control) dataset as well as two mock communities. Based on the clinical sample, we found that NG-Tax1 and NG-Tax2 were strikingly similar in terms of the number of reads/OTUs/ASVs, number of singletons, number of unclassified reads/OTUs/ASVs at the genus level, and number of phyla and genera. This abundance table characteristics were reflected in the results of the beta-diversity analysis, where NG-Tax1 and NG-Tax2 clustered together based on the genera relative abundance. In both versions of NG-Tax, the same genera were indicated as nominally significantly different, and the FC was almost the same. While output of both NG-Tax versions largely overlapped, output varied greatly compared to the other pipelines (QIIME 1, QIIME2, mothur) in terms of, amongst others, the number of singletons, number of unclassified reads/OTUs/ASVs at the genus level and number of genera. Consequently, we showed that only 40%of genera overlap between all the pipelines. The percentage increased to 70% when applying a 10% prevalence cut-off, thereby only comparing genera with RA > 0.1%. The beta-diversity results indicated that, although the samples cluster better according to sample ID than bioinformatics pipelines, all pipelines detected different patterns of microbial composition (Bray-Curtis) and structure (Jaccard), where QIIME1 diverged the most from the other pipelines. In terms of taxonomy, the most abundant genera across the pipelines differed significantly between the pipelines. More importantly, the conclusions of the case-control comparison varied; out of 10 unique genera showing a case-control difference, only one overlapped between all 5 pipelines. Pipelines differed not only in the number of genera showing a case-control difference, but also in the magnitude and even direction of this effect. Overall, the results indicate a clear lack of consistency across the pipelines.

Based on the MCs, we found that QIIME1 and mothur overestimated genus richness, where NG-Tax1, NG-Tax2 and QIIME2 approached the expected genus richness. Beta-diversity analyses indicated that the pipelines differed in representing expected microbial composition and structure, with NG-Tax1 and NG-Tax2 clustering together. Furthermore, correlation analysis between observed and expected MC indicated that, of all pipelines, QIIME2 came closest to the expected microbiome composition. Comparative analysis of individual genera showed that the average relative abundance of specific taxa varied depending on the bioinformatic pipeline. Overall, MC-based results confirmed that the output of pipelines differed in terms of microbiome composition and structure. These results show how the choice of bioinformatic pipeline not only impacts the analysis of 16S rRNA gene sequencing data but also the downstream association results.

### Pipeline characteristics

QIIME1 yielded different results compared to its successor QIIME2 and the other pipelines, mainly regarding the highest number of total and median reads per sample, (unclassified) OTUs and prevalence-filtered genera. Since January 2018, QIIME1 is not supported anymore by developers and has been replaced by QIIME2. This suggests that if data processed using QIIME1 would be reanalysed with QIIME2 or another pipeline, it would yield different results. Furthermore, we observed that QIIME1 yielded the highest number of unique taxa [6, 25]. MC-based results suggested that QIIME1 (and mothur) overrepresented bacterial richness. Thus, in agreement with Prodan et al. (2020), our advice is that for biological conclusions based on alpha-diversity, QIIME1 users should switch to another pipeline or at least confirm their results with another pipeline [6]. For users interested in low frequency taxa, our study showed that QIIME1 and mothur are most appropriate, as they detected more low abundant genera (abundance <0.01%) compared to QIIME2, NG-Tax1 and NG-Tax2 (with NG-Tax being stricter than QIIME2); however, researchers should take into account that this comes at the costs of having a higher number of spurious taxa.

There is dispute in the research community regarding the matter of keeping or removing singletons, and on the best method to remove them. By default, mothur and QIIME2 keep the singletons (69.5% of total OTU/ASVs compared to 0.14% in QIIME2). Both pipelines have different ways of dealing with singletons [19, 21], where mothur yielded highest percentage of singletons. Many of these reads might be noise [43]. Indeed, based on the MCs, we saw that singletons might explain a large number (65% in case of MC3, 40% in case of MC4) of spurious genera (data not shown). However, effects on relative abundance were limited, since singletons accounted for only 0.64% of total reads (for the NeuroIMAGE dataset). Based on these results, we suggest to remove singletons even with the pipelines that suggest keeping them. In addition to the effects on the structure (presence/absence of genera), very low frequency values pose a great challenge for statistical analysis. This is especially relevant if data are analysed at the OTU/ASV level.

This is the first time the output (relative abundance table) of the five pipelines is used together to detect case-control differences and evaluate their consistency and stability in a common statistical framework. Other researchers compared some of these pipelines, and findings partly overlap with ours. For instance, Ducarmon et al. (2020) compared NG-Tax1 and QIIME2 and concluded that the pipelines showed different results in terms of richness [13]. In concordance with our study, NG-Tax1 accurately retrieved richness at the genus level. However, QIIME2 overestimated genus-based richness, whereas in our paper it approached the expected richness in MCs. Furthermore, we observed that the choice of pipeline influenced the analyses of bacterial composition and structure, whereas in the analysis reported by Ducarmon et al. (2020), diversity and compositional profiles were comparable.

With regard to the MCs, in Ducarmon et al. (2020), QIIME2 failed to classify *Salmonella*, and NG-Tax1 detected *Salmonella*, whereas in our study, none of the pipelines detected this genus. This could be due to the difference in the expected RA. In our case, it was 1.2% for MC3 and 2.5% for MC4. For Ducarmon et al. (2020), it was approximately 9%. When looking closer at QIIME2 performance, Almeida et al. (2018) suggested QIIME2 as an optimal pipeline for 16S rRNA gene profiling based on the lowest distance between the expected and observed sample compositions based on synthetic, simulated datasets, and based on the best recall and precision [44]. We observed similar results, where correlations between expected and observed MC composition where highest for QIIME2. In addition to that, according to Prodan et al. (2020), DADA2 (we used QIIME2 with the DADA2 option as a denoising algorithm) offered the best sensitivity, detecting all 22 true ASVs present in their MC [6]. Moreover, our results agree with those of Allali et al. (2017), where DADA2 resulted in lower numbers of ASVs when compared to the number of OTUs of QIIME1 [25] and mothur (this paper); however, this was not seen when comparing QIIME2 to NG-Tax1 and NG-Tax2, suggesting that NG-Tax is even more strict then QIIME2 in terms of quality control settings (e.g., abundance threshold). Altogether, based on our results and existing comparisons, QIIME2 (or DADA2) is a highly recommended pipeline for microbiome research.

Studies investigating differences between bioinformatics pipelines have so far focussed on general characteristics of the OTUs/ASVs/reads such as richness, diversity and microbial compositional profiles rather than the biological conclusions to be drawn from comparing these characteristics e.g., between clinically relevant groups [6, 13, 14]. One study investigating if the same biological conclusions could be reached using different pipelines was Allali et al. (2017), based on data from chicken cecum microbiome (vaccinated, prebiotic treated, control group). They tested different settings of QIIME1, UPARSE and DADA2 and concluded that they could discriminate samples by treatment, despite differences in diversity and abundance, leading to similar biological conclusions [25]. Allali et al. (2017) based their conclusion on beta-diversity rather than a comparative analysis of individual genera (as presented in the current paper). However, they reported differences in RA of specific genera between pipelines, suggesting that also in their data different pipelines resulted in different lists of genera discriminating between treatments. In our study, MC analysis helped to interpret clinical data. The results (e.g., beta-diversity) showed that the MC-based analysis does not necessarily reflect the real dataset as the complexity of a real microbiota sample is much larger. This underlines the importance of deciding which pipeline best serves the analysis of your dataset based on how this pipeline performs on real data as well as MCs.

### Limitations and open questions

Our results should be viewed in the context of some limitations. Our study was limited by a small sample size (N=90), but taking into consideration that this is a crossover study the sample size should be sufficient to detect the differences in the output produced by each of the pipelines and how these differences affect the downstream statistical analysis. Nevertheless, since microbiome data is notoriously diverse and sensitive to protocol and technical variations [45, 46], the effect of datasets with different designs should be investigated. Another limitation of this study was the use of nominal (and standard) statistical significance cut-off (p<0.05) as a measure of statistical difference. Considering the number of tested genera, several false positives could be expected. Although a corrected p-value is considered a better measure of success, the case-control study may not contain large enough differences or enough statistical power to properly classify the differences between groups as statistically significant. Given the aim of this paper, establishing the true (biological) difference between groups is not evaluated and comes second to the difference in observed effects brought in by the choice of the bioinformatic pipeline, which is why nominal significance was sufficient to select multiple taxa (showing different RA and p-values across pipelines) and evaluate the effect on analysis. Lastly, the number of ASVs/OTUs varied considerably between pipelines, which can result in differences in FC magnitude, as seen for example in case-control ratio differences between QIIME1 and QIIME2 on the *Clostridiales vadin* genus group. A different direction of FC could be driven by a differential effect of filtering/denoising steps per group, potentially driven by a larger number of sequencing artefacts in either of them. Future research should focus on more technical aspects of bioinformatics pipelines comparisons, to identify what exactly drives such differences.

## 5. Conclusion

Our results indicate that a choice of bioinformatic pipeline has not only an impact on the analysis of 16S rRNA gene sequencing data but also the case-control comparison results. This means that the choice of pipeline can influence the list of significantly different genera between study groups. Thus, we underscore a significant limiting factor in current microbiome research: the lack of consistency between study results and how this limits their comparability and the validity of conclusions to be drawn from them.

Based on our results we recommend i) using QIIME1 and Mothur to researchers that are interested in rare and/or low-abundant taxa, ii) using NG-Tax1 or NG-Tax2 when favouring strict artefact filtering, iii) using QIIME2 when looking for a balance between the two abovementioned points, and iv) using at least two pipelines to assess the stability of results.

We would like to point out that the field still needs to develop “best practice” for microbiome analysis and apply it consensually across studies, before we can have a deeper understanding of the gut microbiota’s contribution to human health and disease. With our current work, we hope to contribute to the gut microbiota research field and make other researchers aware of the strengths and limitations of their choice of bioinformatic pipeline in terms of influencing the results of case-control studies with 16S rRNA marker gene sequencing data.

## Declarations

### Ethics approval and consent to participate

The study was approved by the Institutional Review Board at Radboud University Medical Centre, Nijmegen, The Netherlands (registration number 2012/542; NL nr.: 41950.091.12). An informed written consent was obtained from all participants and/or their parents prior to the sample and data collections.

### Data Availability

The data underlying this article will be shared on reasonable request to the corresponding author.

### Competing interests

The authors declare that they have no competing interests.

### Funding

This project was supported by the European Union’s Horizon 2020 research and innovation programme under the Marie Sklodowska-Curie [grant agreement number 643051]. The funding body had no role in the preparation of the manuscript. The research leading to these results also received funding from the European Community’s Horizon 2020 research and innovation programme under project no. 728018 (Eat2BeNICE) and 667302 (CoCA).

## Acknowledgements

The authors would like to thank Prokopis Konstanti, Ward de Witte, Fini de Gruyter, QIIME2 forum (https://forum.qiime2.org/) and mothur forum (https://forum.mothur.org/) for helpful discussions.

## Supplementary Figures

**Figure S1.** Results of Jaccard similarity index. *A)* Principal Coordinates Analysis (PCoA) plots with the percentage explained variance by the principal coordinates. *B)* Canonical Analysis of Principal coordinates (CAP) ordination plot reveals structure in microbial communities associated with bioinformatics pipelines. *C)* TukeyHSD, a pairwise comparison of group mean dispersions, revealed that the intra-sample variation is quite similar across pipelines, except for QIIME1.

**Figure S2.** Scatter plot representing a correlation of the PCS with the relative abundance (A) and prevalence (B) based on the 10 genera showing nominally significant differences (p< 0.05) between cases and controls in at least one pipeline (Table 2).

**Figure S3.** Correlation matrix of Spearman correlation coefficient values between observed mock community (OBS MC) composition as a result of five different bioinformatics pipelines (NG-Tax1, NG-Tax2, QIIME1, QIIME2 and mothur) and corresponding expected mock composition (EXP MC), Mock_3 (A) and Mock_4 (B). The results are based on genera only present in EXP MCs. The observed values represent statistically significant correlations (P<0.05).

**Figure S4.**Interactive heatmap of the expected (EXP) and observed (OBS) MC3 based on all genera (N=36) present in EXP MC. The rows of the matrix are ordered to highlight patterns by using default settings.

**Figure S5.**Interactive heatmap of the expected (EXP) and observed (OBS) MC4 based on all genera present (N=36) in EXP MC. The rows of the matrix are ordered to highlight patterns by using default settings.

## Supplementary Tables

**Table S1.**Results of pairwise PERMANOVA.

**Table S2.**Descriptive statistics of 16 most abundant genera.

**Table S3.**Descriptive statistics of 10 genera shown in Table 2.

**Table S4.**Number of genera based on observed and expected (EXP) MC.

## Notes

### Competing Interest Statement

The authors have declared no competing interest.

